# Multiplexing Mutation Rate Assessment: Determining Pathogenicity of Msh2 Variants in *S. cerevisiae*

**DOI:** 10.1101/2020.10.05.325902

**Authors:** Anja R. Ollodart, Chiann-Ling C. Yeh, Aaron W. Miller, Brian H. Shirts, Adam S. Gordon, Maitreya J. Dunham

**Affiliations:** Molecular Cellular Biology Program, University of Washington, Seattle, WA; Genome Sciences Department, University of Washington, Seattle, WA; Zymergen, Emeryville, CA; Department of Laboratory Medicine, University of Washington, Seattle, WA; Department of Pharmacology, Northwestern University, Chicago, IL

## Abstract

Despite the fundamental importance of mutation rate as a driving force in evolution and disease risk, common methods to assay mutation rate are time consuming and tedious. Established methods such as fluctuation tests and mutation accumulation experiments are low-throughput and often require significant optimization to ensure accuracy. We established a new method to determine the mutation rate of many strains simultaneously by tracking mutation events in a chemostat continuous culture device and applying deep sequencing to link mutations to alleles of a DNA-repair gene. We applied this method to assay the mutation rate of hundreds of *Saccharomyces cerevisiae* strains carrying mutations in the gene encoding Msh2, a DNA repair enzyme in the mismatch repair pathway (MMR). Loss-of-function (LOF) mutations in *MSH2* are associated with hereditary non-polyposis colorectal cancer (HNPCC), an inherited disorder that increases risk for many different cancers. However, the vast majority of *MSH2* variants found in human populations have insufficient evidence to be classified as either pathogenic or benign. We first benchmarked our method against Luria-Delbrück fluctuation tests using a collection of published *MSH2* variants. Our pooled screen successfully identified previously-characterized non-functional alleles as high mutators. We then created an additional 185 human variants in the yeast ortholog, including both characterized and uncharacterized alleles curated from ClinVar and other clinical testing data. In a set of alleles of known pathogenicity, our assay recapitulated ClinVar’s classification; we then estimated pathogenicity for 157 variants classified as uncertain or conflicting reports of significance. This method is capable of studying the mutation rate of many microbial species and can be applied to problems ranging from the generation of high-fidelity polymerases to measuring the rate of antibiotic resistance emergence.

## Introduction

Mutation rate is the timer for many different error-prone processes: how many cycles of PCR before the polymerase makes a mistake, how long before the bacterial infection becomes resistant to existing medications, or how quickly will DNA damage result in the uncontrolled growth of a cancerous tumor. An example of the latter is germline variants in mismatch repair (MMR) pathway genes which have strong implications for human health and are responsible for the cancer risk syndrome known as hereditary non-polyposis colorectal cancer (HNPCC) (Lynch et al., 2015; Peltomäki, 2016). In patients carrying pathogenic alleles, increased surveillance can detect cancers early, improving treatment outcomes(Gupta et al., 2019). There exist a large number of variants of MMR repair genes in humans, and many are classified as variants of uncertain significance (VUS)(Starita et al., 2017). Functional data found in model organisms can be used to assess potential pathogenicity of these variants(Gordon et al., 2019; Richards et al., 2015). As such, a way to screen variants to determine if they cause a heightened mutation rate, and so therefore may be pathogenic, is needed.

However, existing methods for measuring mutation rate are tedious and not scalable for the challenge of functionally testing hundreds or thousands of VUS. Methods to study mutation rate all have their benefits and detriments(Foster, 2006). In microbial systems, Luria-Delbrück fluctuation tests and mutation accumulation lines are two of the most used(Luria and Delbrück, 1943; Lynch et al., 2008). All of these methods require initiating populations with single clones to determine the effect of a mutation, which limits the ability to multiplex experiments. An alternative method is measuring sensitivity to 6-thioguanine or N-Methyl-N’-Nitro-N-Nitrosoguanidine, which correlates with functionality of subunits of the MMR complex including Msh2 and Mlh1(Bouvet et al., 2019; Drost et al., 2013; Houlleberghs et al., 2017; Jia et al., 2020). Genome editing methods such as CRISPR-Cas9 provide a convenient way to introduce variants into human cells, where signatures of MMR deficiency can then be tracked(Rath et al., 2019). Another alternative is cell free systems, which allow for using the human protein in an assay that checks for ability to repair DNA(Drost et al., 2010, 2012, 2019). While this provides an easy way to see all mutations caused by errors in replication, each variant must be expressed and purified separately and this strategy is not amenable to being pooled. Also, while these systems work well for MMR complex proteins, they do not generalize to all sources of mutation rate variability. Computational strategies theoretically could scale to all possible sites in all proteins of interest, and have been demonstrated to be good predictors of destabilizing variants of two MMR proteins tested in human colorectal cell lines (Abildgaard et al., 2019; Nielsen et al., 2017; Stein et al., 2019). However, they still require further validation. To address some of these problems, we wanted to generate a new experimental protocol to do multiplexed, direct assessment of mutation rate that was amenable to any molecular pathways for which mutations can be a read out in an easy to culture and genetically tractable organism, *Saccharomyces cerevisiae*.

In addition to ease of use, yeast is a good model system to study effects on MMR because much of the sequence and function of the pathway is conserved between humans and yeast(Boiteux and Jinks-Robertson, 2013). Many discoveries about MMR originate in studies of *S. cerevisiae*, as the MMR complex is more highly conserved with its human orthologs than that of *E.coli* (Strand et al., 1993, 1995). In addition to general biology of the MMR complex, yeast has been used to determine the mutation rate of MMR alleles using traditional fluctuation assays, mutation accumulation lines, qualitative patch assays, and fluorescence-based assays (Demogines et al., 2008; Drotschmann et al., 1999a; Gammie et al., 2007; Lang and Murray, 2008; Lang et al., 2013; Martinez and Kolodner, 2010; Shor et al., 2019). These assays all test the full functionality of Msh2, however none of them allow for pooled assessment of many alleles simultaneously. Although medium-throughput assays exist that take advantage of automated liquid-handling systems(Gou et al., 2019), these still require considerable effort if studying the effect on mutation rate of many different alleles and require keeping clonal populations.

In an attempt to improve on the scaling problems of other methods for measuring mutation rate, we have developed a chemostat-based assay that utilizes pools of variants to assay hundreds of alleles simultaneously. The chemostat is a continuous culture device which maintains a constant population size over time. The first use of the chemostat to determine mutation rate was over half a century ago (Fox, 1955; Kubitschek and Bendigkeit, 1964; Novick and Szilard, 1950; Paquin and Adams, 1983). The continuous dilution in the chemostat means an increase in the frequency of a neutral mutation is a result of *de novo* mutation, as opposed to other methods, where such an increase could be explained by both *de novo* mutation and exponential growth. If the neutral mutation is also selectable, such as with some types of drug resistance (e.g. canavanine resistance in yeast), the mutation rate can be calculated by simply tracking the frequency of resistance over time. Combining this very old technique with next-generation sequencing allows for the pooling and therefore high-throughput study of alleles. We applied this new method to study mutation rate differences caused by variants in *MSH2*, a gene that is associated with HNPCC. Msh2 is a part of the MMR complex, which in combination with Msh3, Msh6, Mlh1, and Pms2 binds and fixes small mismatches and indels (Boiteux and Jinks-Robertson, 2013). Msh2 is an integral part of the recognition complex (Edelbrock et al., 2013). We completed a proof of principle with previously published variants of *MSH2*, and found that the pooled assay recapitulated the results of traditional Luria-Delbruck fluctuation tests, qualitative patch assays, and yeast two-hybrid assays (Gammie et al., 2007). We then assayed an additional 185 *MSH2* missense variants curated from ClinVar, a public repository of clinical variant interpretations derived from diagnostic genetic testing. To do so, we recreated these variants in the homologous sites of yeast *MSH2*, barcoded them along with control wt clones, and measured their mutation rates in a pooled format. Of the 28 variants of known pathogenicity, 100% recapitulated the functional consequence implied by previous clinical interpretation. We then examined 157 VUS from ClinVar and identified 50 variants with significantly different mutation rates from WT as measured by our assay. In addition to ClinVar classifications, data were also compared to tumor sequencing in cancer patients (Shirts et al., 2018); of the 25 VUS for which tumor data was available, 64% had clinical findings that were consistent with our functional data. These data taken together show that our new multiplexed mutation rate assay is an accurate and scalable assay to study the mutation rate of many strains in a pooled format.

## Results

### A new method for multiplex mutation rate assessment

The chemostat is a continuous culture device which matches the growth rate of an organism to the dilution rate, stabilizing population size and environmental conditions throughout an experiment. Many ways to study mutation rate take advantage of drugs for which WT organisms are sensitive but a loss-of-function mutation causes resistance, which makes it straightforward to track mutation frequency(Boeke et al., 1987; Whelan et al., 1979). However, to determine rate, one must know the number of generations that have elapsed since the mutational event, which is difficult in batch culture. Luria-Delbrück fluctuation assays and mutation accumulation lines use different tactics to determine the generational time. In continuous culture, since the population size stays stable, an increase in resistance isn’t due to an increase in a lineage, as long as certain underlying assumptions are met, as described in the following section. In the assay we have developed, outlined in Fig 1, we can track many lineages in a pooled manner to determine all of their mutation rates at once. We do this by keeping track of *de novo* mutational events on selective media containing a drug, while controlling for any changes in overall population size by monitoring growth on non-selective media. Next-generation sequencing of the plasmid recovered from both the non-selective and selective media allows us to track the various lineages over time. This assay is amenable to both barcoded and unbarcoded libraries. With unbarcoded libraries, we use shotgun sequencing of the allele isolated from the plasmid, using the mutation within the gene itself as a way to track the variant over the course of the experiment. In barcoded libraries, the barcode and variant are first linked using long read sequencing, after which amplicon sequencing of just the barcode reveals the frequency of each variant at each time point. Our method is amenable to both types of analysis to make it more generalizable. In both cases, the increase in resistance frequency over time for all lineages can be calculated, giving us their mutation rates. Our first application of this method utilizes *Saccharomyces cerevisiae* and focuses on variants of Msh2, a clinically relevant DNA repair enzyme. However, this assay is amenable to the study of any microbial strains which can be cultured within the chemostat and any molecular pathway that yields a neutral selectable mutation as a phenotypic readout.

**Fig 1.**
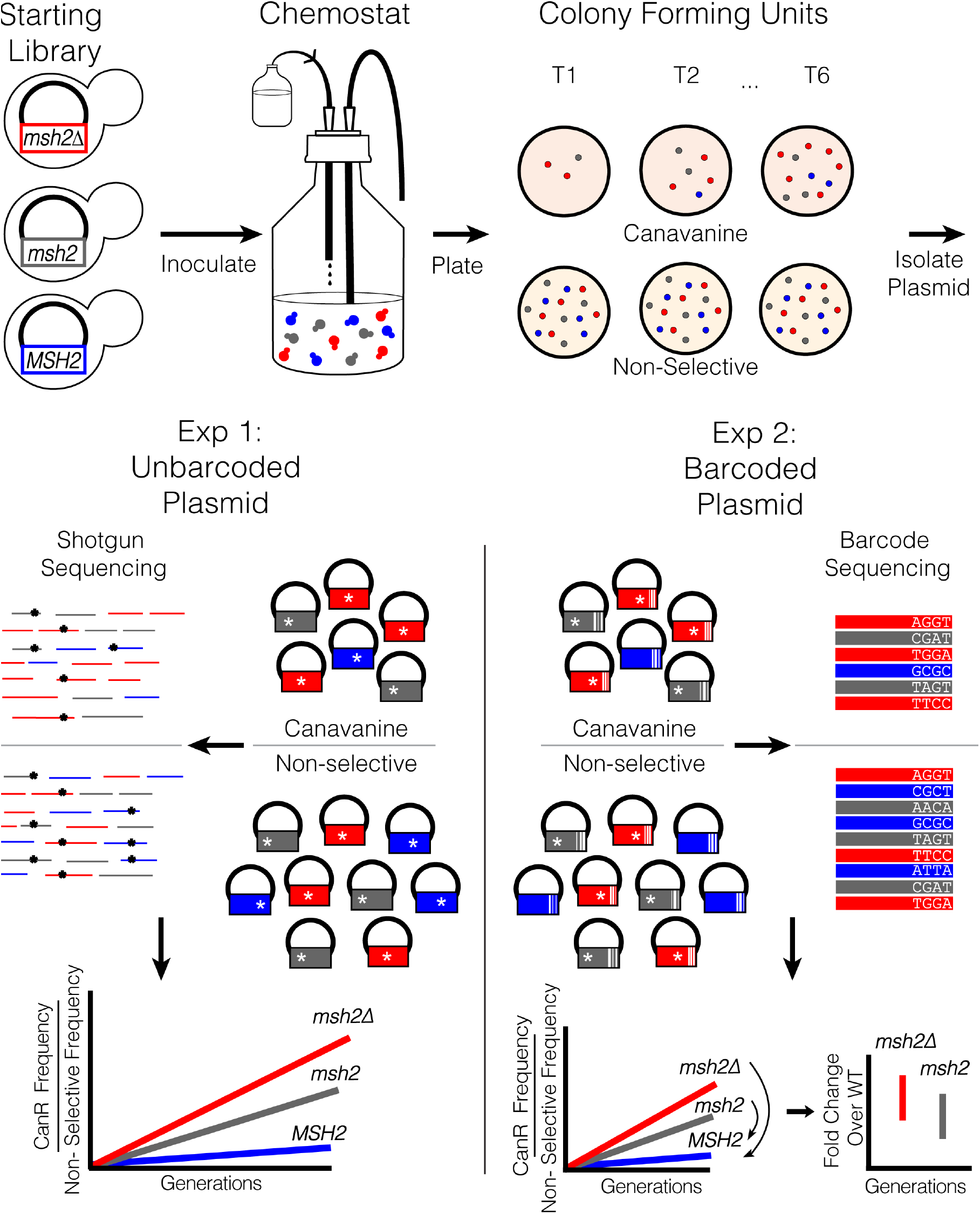
A schematic outlining the multiplexed mutation rate method. A pool of alleles is inoculated into a chemostat. Samples are plated onto non-selective media or media containing canavanine to select for LOF mutations in *CAN1*. Plasmid is isolated from the canavanine selected plates as well as from the non-selective pool. The assay can handle both unbarcoded and barcoded plasmids, using shot gun or barcode sequencing respectively. In both cases the frequency of the allele on selective canavanine media is divided by presence in the total pool and tracked over time to generate the mutation rate. With barcoded plasmids, barcoded WT can be used to determine the fold change of variants against an internal control.

### Conditions required for mutation rate assessment in the chemostat

To accurately measure mutation rate in the chemostat some underlying criteria must be satisfied. First, the readout of mutation rate should be neutral in fitness. Second, resistance should accumulate linearly over the assay period indicating no spontaneous non-neutral mutations arose and reached a detectable frequency. Lastly, there should be no inherent fitness effect due to the variants we seek to characterize. If these underlying criteria are unmet, then the link between an increase in the drug resistance marker and *de novo* mutation is broken.

To address the first assumption, we used resistance to canavanine as a read out. Loss-of-function mutations in *CAN1*, an arginine transporter, prevent uptake of the toxic arginine analog canavanine since the transporter is non-functional. Canavanine has been successfully used to assay mutation rate in yeast previously(Lang and Murray, 2008; Paquin and Adams, 1983) and has a minimal 1.015% fitness benefit in glucose limitation, the conditions used in this study(Gresham et al., 2008).

To determine the timeframe over which we could observe linear accumulation of resistance, we assayed the mutation rate of *msh2Δ* strains containing either WT *MSH2* (*MSH2*) or pRS413 (*msh2Δ*) using the same conditions as future pooled experiments. Previous work has shown complementation by plasmid-borne WT *MSH2*, and that variants that abrogate activity elevate the mutation rate in yeast (Strand et al., 1993; Drotschmann et al., 1999b; Gammie et al., 2007). Chemostats were individually inoculated with *MSH2* and *msh2Δ* strains, and samples were plated every 24 hours to determine which timepoints correspond to the range for linear accumulation of Can^R^ mutants [Supp fig 1]. We found that between 12 and 45 generations, resistance to canavanine accumulates at very similar rates as stated in previous literature (Table 1) (Gammie et al., 2007). The lag in linear accumulation can be explained by a requirement for cells to reach steady state, at which point they are growing and being diluted at the same rate (reviewed in (Gresham and Dunham, 2014). After 45 generations, selection on adaptive mutations is likely the cause of the non-linear increase(Paquin and Adams, 1983). From this, we determined that all future experimental timepoints must be taken between 12 to 45 generations to accurately determine the mutation rate.

**TABLE 1.** Mutation Rates of Control Pure Cultures and Pools.

| Relevant Genotype | CanR rate<sup>a</sup> | Fold Induction<br>CAN |
| --- | --- | --- |
| *MSH2* | $1.52 \times 10^{-7} \pm 0.07$ | 1 |
| *msh2Δ* | $49.44 \times 10^{-7} \pm 2.6$ | 29 |
| Unbarcoded pool | $21.62 \times 10^{-7} \pm 2.53$ | 13 |
| Barcoded Pool | $39.6 \times 10^{-7} \pm 2.1$ | 26 |

a. Rate, mutations per generation

If we are to multiplex this assay, null-like *msh2* variants should not have a large fitness effect, otherwise we cannot determine the difference between a *de novo* mutation and expansion or contraction of a resistant lineage. In a head-to-head competition between WT and *msh2Δ*, we found a 1.097% fitness defect of *msh2Δ* over the short course of our experiment (sup fig 1). This means we will likely slightly underestimate mutation rate of high mutators. However, by correcting our mutation rates by the relative population frequency of each variant, we can mitigate the effects of both the strain manipulation and resistance to canavanine.

### Pools of alleles accumulate mutations at expected rates

Mutation rate can vary even under very similar conditions, and thus multiple replicate assays must be done to obtain accurate measurements. We created a pooled assay to easily increase the number of replicates for each individual mutation rate assessment. We first established a proof of principle assay with 46 previously published alleles of *MSH2*(Gammie et al., 2007) (fig 2). We pooled equal amounts of strains carrying each allele, inoculated aliquots of the pool into 4 independent 200ml chemostats, and from each collected 6 samples on selective and nonselective media over 50 generations, as determined from the control experiments described above. Plasmid was isolated from all samples, plasmid-borne *MSH2* alleles were amplified by PCR and subjected to shotgun short read sequencing, and the frequency of each allele from the canavanine resistant pool was normalized by the frequency in the non-selective pool and converted into colony counts (see methods).

**Fig 2.**
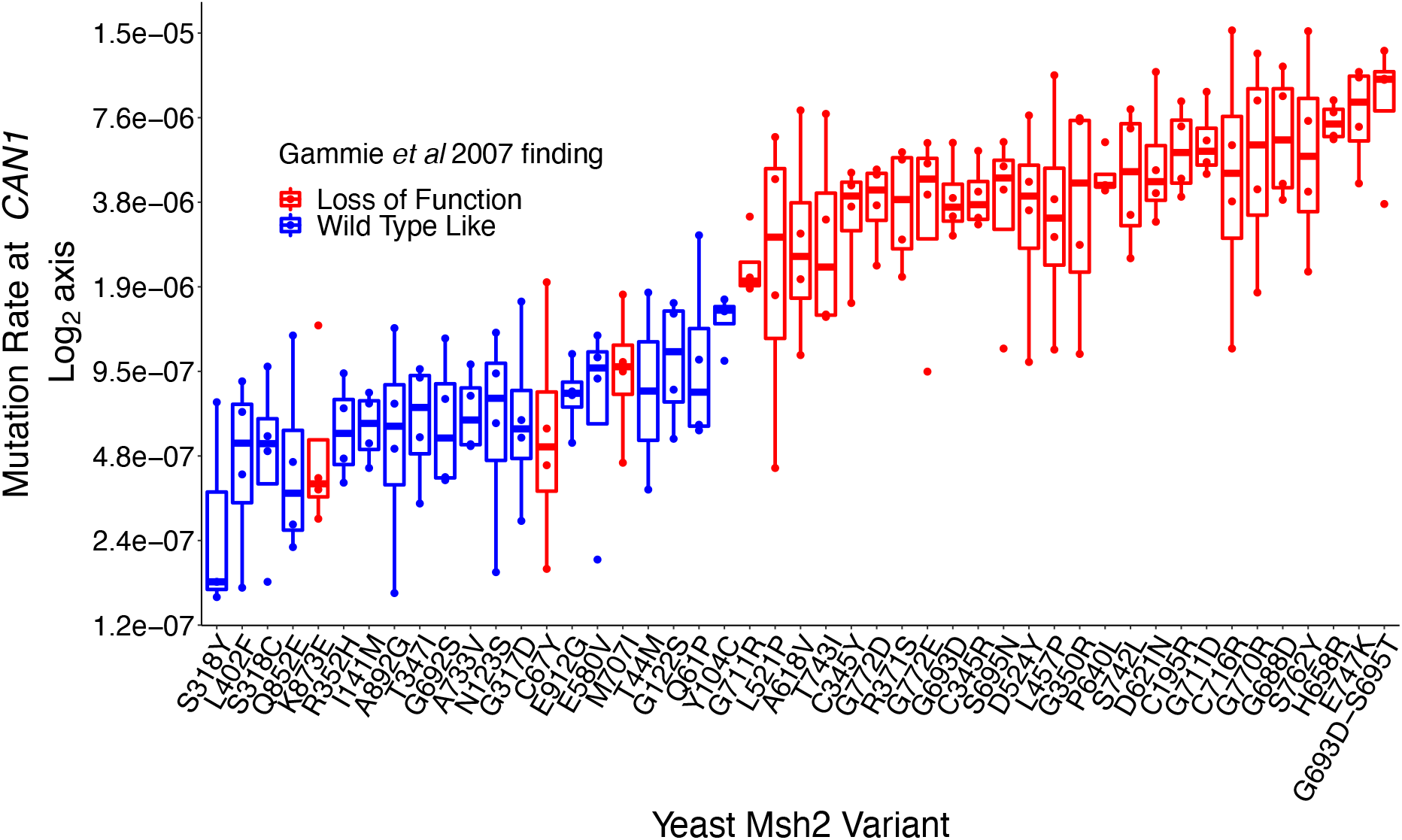
Calculation of mutation rates of indicated alleles using multiplexed mutation rate assessment. Mutation rate (CanR per generation) of previously studied alleles, colored by the phenotype found in Gammie *et al* 2007. Mutation rates are plotted on a log_2_ axis and points represent measurements from separate chemostats.

The average mutation rate of this pool is 13X over wild type (Table 1), which corresponds with what is expected in a pool containing both loss-of-function and WT-like alleles. In Figure 2, the mutation rates of each allele as measured at *CAN1* are plotted. We compared the results to the phenotype found in previous work using qualitative patch assays, Luria-Delbrück fluctuation tests, and yeast two-hybrid assays (Gammie et al., 2007). With the exception of 3 alleles (K873E, C67Y, and M707I), alleles previously described as WT-like all grouped together, as did the loss-of-function alleles. C67Y was classified as loss-of-function due to a lack of subunit interaction and a qualitative patch assay in previous work. This lack of subunit interaction may not be reflected in our mutation rate assay and perhaps explains the lack of correspondence to the qualitative measurement. K873E and M707I both showed a LOF phenotype measured at *CAN1* but were found to be WT-like when testing for dinucleotide instability. These alleles exist at the edge of the classification between loss of function and WT-like in the compared study and could potentially explain why the results are discordant. These data show a grouping of the high mutators and low mutators, indicating that our new method can largely replicate the results of previous efforts to measure allele-specific effects on mutation rate.

### Simultaneous measurement of mutation rate among 2000 different lineages

To determine the limit on the number of alleles that can be assayed at once, additional alleles of Msh2 were curated from the clinical database ClinVar. These human Msh2 variants were mapped onto the yeast *MSH2* gene; only sites with a residue conserved between both orthologs were considered. 28 alleles with known pathogenicity were included as well as 216 variants of uncertain significance. Alleles in the Walker A ATPase domain and the linker region, which are more highly conserved between humans and yeast, were given precedence(Gammie et al., 2007). Alleles were barcoded a median of 5 times (supp fig7) and the barcode and variant were associated with long read Pacific Biosciences sequencing. Of the 244 alleles synthesized, 191 variants covered by 1261 barcodes were able to be assayed. In addition, 737 barcodes were associated with WT, giving a robust internal control (Fig 3A). The 53 variants not assayed were due to low barcode coverage of the variant in the pool or other factors that caused them to fail our quality filters (supp Figs 2-3). While the number of variants that can be assayed is dependent on the composition of the pool, with higher mutators being easier to assay than lower mutators, results from this pool of Msh2 variants indicates that the assay is capable of tracking approximately 2000 barcodes at once.

**Fig 3.**
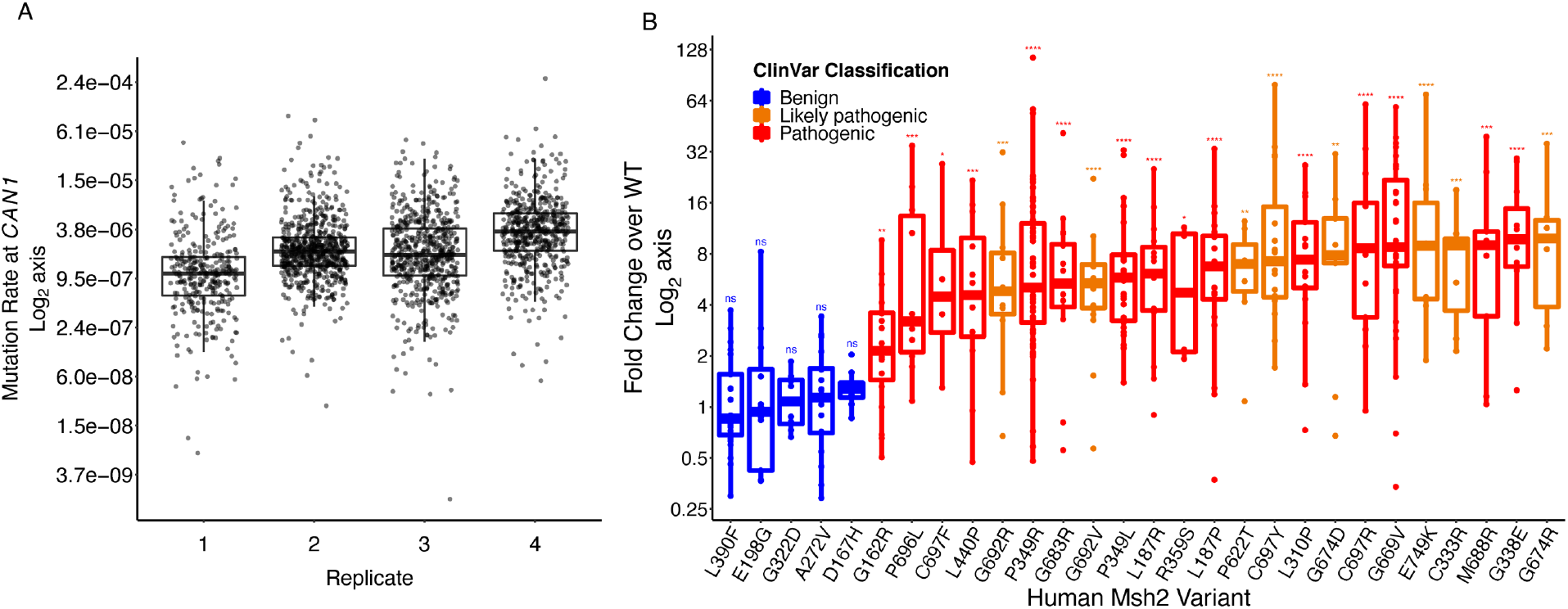
Control alleles of Msh2 in barcoded experiment pool. A. Calculated mutation rates (CanR per generation) of WT barcodes in four replicate experiments plotted on a Log_2_ axis. B. Fold change over WT plotted on a Log_2_ axis, colored by pathogenicity classification in ClinVar. Points represents individual barcoded measurements from the four replicate experiments. Significance determined by comparing variants to the WT barcodes by a Wilcoxan Rank-Sum test with the Benjamini-Hochberg correction for multiple hypothesis testing. *<0.05 ** <0.01 ***<0.001 ****<0.0001

### Internal WT barcodes can identify differences in mutation rate among alleles

The addition of barcodes, while requiring additional work and cost, can provide internal controls to the pooled mutation rate assay, as each genotype can be tracked in multiple independent lineages. The cumulative mutation rate of the barcoded pool containing novel variants of Msh2 was 3.96 × 10^−6^ Can^R^/generation, a 26-fold increase over WT (Table 1). The WT barcodes in this assay showed a median mutation rate between 1.09 x10^−6^ – 3.59 x 10^−6^ – approximately 10-fold higher than the expected rate (Fig 3A). While the source of this global increase in mutation rate is unknown, it can be controlled for by the internal WT barcodes. All variant barcodes in the pool were compared to the internal median WT mutation rate of that pool, to calculate fold-change over WT for each variant. Inclusion of barcoded WT allows for a robust internal control for mutation rate assessment, which mitigates spurious sources of increased mutation.

### Allele-specific mutation rates recapitulate ClinVar pathogenicity classifications

To determine if the variants in the barcoded pool recapitulated clinical variant interpretations reported from diagnostic testing, we first looked at variants with known pathogenicity scores in ClinVar. The control variants used have a review status of 2 or higher indicating strong evidence for variant classification (Fig 3B). Variants that shared the same ClinVar classification were found to have similar mutation rates. All 5 benign variants showed no significant difference from WT. 23 out of 23 pathogenic and likely pathogenic alleles showed a significant increase in mutation rate compared to WT. Due to the limited number of previously characterized variants in the pool, it’s difficult to determine true sensitivity and specificity scores; however, these data lend confidence to our ability to bin variants into pathogenic or benign categories. Based on the results of known variants, we have created four bins: variants which do not differ significantly from WT are potentially benign (1). Those with values ranging between 1.3X −1.4x over WT are likely intermediate (2). Those which are significantly higher than WT are potentially pathogenic (3). The lowest fold change which showed a significant difference from WT was 1.4X, so we classified those which are above this threshold but do not reach significance as possibly pathogenic (4). We were heartened to see our assay recapitulates previous clinical interpretations, and that the use of control alleles allowed for the generation of bins to provide information on the VUS assayed.

### Estimating pathogenicity of variants of uncertain significance

We were able to assay 157 SNP variants of uncertain significance in this assay. 50/157 showed a significant difference in mutation rate in comparison to WT, and were classified as potentially pathogenic. The mutation rates of these 50 variants are in table 2. A summary of the fold changes of all variants is found in supplemental table 1. We found 1 variant, K449N, had a significantly lower mutation rate than WT. The mechanism of this lowered mutation rate and whether it is a biologically relevant is unknown and could provide an interesting point of study if confirmed. The 50 VUS which showed a significant increase in mutation rate ranged from 1.39 fold over WT to 13 fold over WT. No alleles assayed had a full loss-of-function phenotype–which would be characterized by a 30 fold increase in mutation rate. This may reflect the dynamic range of the assay, or it may reflect that no true loss-of-function alleles existed in the data set. For future experiments, it may be wise to include barcoded deletion strains at a low frequency. Taken in total, this data set provides evidence of pathogenicity for an additional 157 VUS of *MSH2*. 83 will be classified as potentially benign, 7 intermediate, 17 possibly pathogenic, and 50 potentially pathogenic. While additional study would be required before these classifications could inform clinical diagnosis, these data represent a first indication of the effects of these mutations on function, and could be used as a line of evidence according to ACMG criteria(Richards et al., 2015).

**Table 2.** Mutation rates of significantly different alleles.

| Human Genotype | Yeast Genotype | Count<sup>a</sup> | Fold Induction CAN<sup>b</sup> | IQR<sup>c</sup> | Sig<sup>d</sup> |
| --- | --- | --- | --- | --- | --- |
| K449N | K466N | 10 | 0.56 | 0.28 | *** |
| G761R | G780R | 31 | 1.39 | 0.96 | *** |
| M672R | M691R | 32 | 1.61 | 3.37 | **** |
| R711Q | R730Q | 25 | 1.62 | 2.63 | **** |
| P349A | P361A | 61 | 1.79 | 2.03 | * |
| V695L | V714L | 12 | 2.08 | 1.77 | **** |
| Q681E | Q700E | 20 | 2.09 | 2.69 | **** |
| R359K | R371K | 7 | 2.25 | 1.26 | **** |
| M672V | M691V | 37 | 2.27 | 2.37 | ** |
| H783Y | H802Y | 11 | 2.38 | 1.82 | **** |
| S676P | S695P | 8 | 2.48 | 4.55 | * |
| A609P | A627P | 12 | 2.56 | 3.36 | ** |
| G761V | G780V | 19 | 2.71 | 2.82 | **** |
| A700E | A719E | 9 | 2.78 | 3.79 | *** |
| H783D | H802D | 7 | 2.79 | 4.53 | * |
| G683E | G702E | 54 | 2.88 | 3.62 | **** |
| A689V | A708V | 11 | 3.17 | 2.58 | **** |
| P696S | P715S | 5 | 3.27 | 1.66 | ** |
| R524H | R542H | 9 | 3.49 | 5.85 | ** |
| R524C | R542C | 13 | 3.61 | 7.10 | **** |
| K675E | K694E | 8 | 3.66 | 7.49 | ** |
| T724M | T743M | 34 | 3.83 | 3.51 | * |
| P622Q | P640Q | 5 | 3.95 | 2.00 | *** |
| P622A | P640A | 9 | 4.12 | 3.17 | ** |
| S676L | S695L | 5 | 4.18 | 1.43 | *** |
| R524L | R542L | 12 | 4.18 | 2.22 | **** |
| Y43D | Y43D | 7 | 4.31 | 2.16 | ** |
| E643K | E662K | 24 | 4.41 | 5.92 | ** |
| G669R | G688R | 13 | 4.90 | 2.62 | **** |
| C693R | C712R | 13 | 5.00 | 10.95 | ** |
| T668P | T687P | 17 | 5.03 | 11.27 | ** |
| G692E | G711E | 14 | 5.04 | 8.82 | **** |
| T724R | T743R | 18 | 5.76 | 6.85 | ** |
| G683W | G702W | 18 | 5.81 | 6.00 | ** |
| G827R | G855R | 34 | 5.99 | 8.72 | ** |

**Table 2** **(Continued)**
| Human<br>Genotype | Yeast<br>Genotype | Count<sup>a</sup> | Fold<br>Induction<br>CAN<sup>b</sup> | IQR | Sig<sup>d</sup> |
| --- | --- | --- | --- | --- | --- |
| Q690E | Q709E | 15 | 6.03 | 8.01 | * |
| A689P | A708P | 9 | 6.27 | 5.14 | *** |
| P670R | P689R | 6 | 6.31 | 6.23 | **** |
| G692W | G711W | 23 | 6.33 | 6.60 | * |
| G669D | G688D | 18 | 6.63 | 11.83 | * |
| G827E | G855E | 11 | 6.95 | 7.11 | **** |
| T677R | T696R | 13 | 7.41 | 4.94 | **** |
| P349S | P361S | 47 | 7.48 | 8.01 | **** |
| G338V | G350V | 14 | 7.98 | 15.14 | **** |
| G669C | G688C | 12 | 8.08 | 5.35 | ** |
| R621Q | R639Q | 4 | 8.29 | 2.07 | *** |
| N671D | N690D | 13 | 8.61 | 6.56 | * |
| R621P | R639P | 13 | 9.28 | 10.78 | **** |
| L310R | L305R | 14 | 9.50 | 13.27 | **** |
| C693Y | C712Y | 4 | 9.70 | 3.71 | **** |
| R680P | R699P | 9 | 13.00 | 17.19 | **** |

a. Count, the number of times a variant was assayed in total
b. For stated genotype, mutation rate (mutations per cell division) was calculated and then compared to the WT mutation rate. Barcode and chemostat replicates are combined to calculate median foldchange.
c. Interquartile range of all barcode and chemostat replicates
d. Significance is calculated by a Wilcoxon Rank-sum test with the Benjamini-Hochberg correction for multiple hypothesis testing. \* <.05 \*\* <.01 \*\*\* <.001 \*\*\*\* <.0001

### Associating variant data with clinical and tumor sequencing phenotypes

In order to more accurately compare clinical data with the outputs of this screen, the fold change calculations were converted to scores. Table 3 contains information on variants which have clinical findings associated with them. Clinical summaries were gathered from data provided to the University of Washington Genetics and Solid Tumors Laboratory. An assessment of whether the clinical information is consistent or inconsistent with functional scores was provided by a board-certified molecular pathologist with expertise in this area (BHS). Clinical evidence was considered consistent with functional data when both suggested the variant was pathogenic or benign regardless of the strength or significance of the data. There are several types of information on *MSH2* that can be gathered from patients, families, and tumors(Thompson et al., 2013; Rañola et al., 2018; Shirts et al., 2018; Li et al., 2020).

**Table 3.**
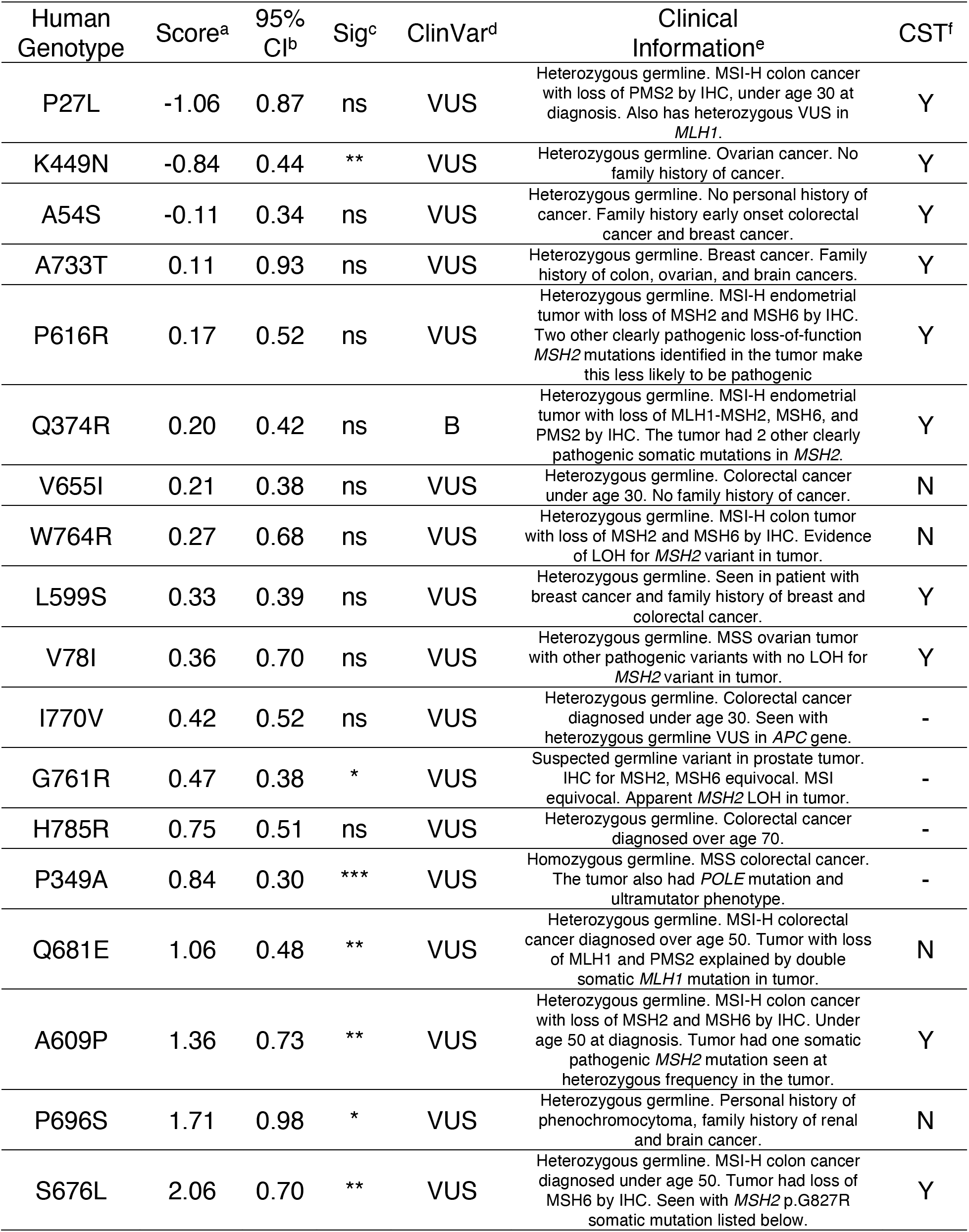
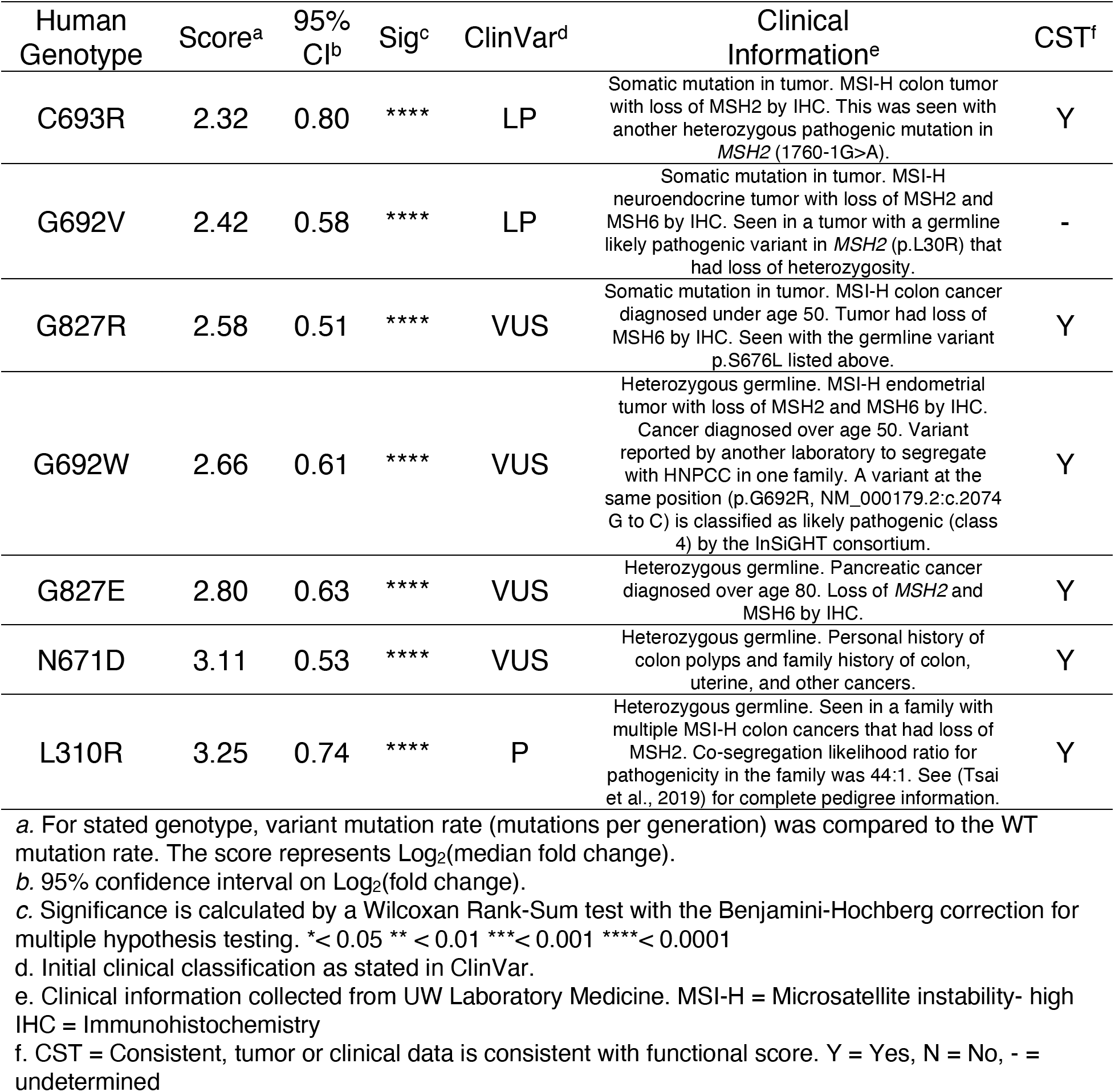
Summary of variants with clinical or tumor data.

Summary of variants with clinical or tumor data
| Human Genotype | Score<sup>a</sup> | 95% CI<sup>b</sup> | Sig<sup>c</sup> | ClinVar<sup>d</sup> | Clinical Information<sup>e</sup> | CST<sup>f</sup> |
| --- | --- | --- | --- | --- | --- | --- |
| P27L | -1.06 | 0.87 | ns | VUS | Heterozygous germline. MSI-H colon cancer with loss of PMS2 by IHC, under age 30 at diagnosis. Also has heterozygous VUS in *MLH1*. | Y |
| K449N | -0.84 | 0.44 | ** | VUS | Heterozygous germline. Ovarian cancer. No family history of cancer. | Y |
| A54S | -0.11 | 0.34 | ns | VUS | Heterozygous germline. No personal history of cancer. Family history early onset colorectal cancer and breast cancer. | Y |
| A733T | 0.11 | 0.93 | ns | VUS | Heterozygous germline. Breast cancer. Family history of colon, ovarian, and brain cancers. | Y |
| P616R | 0.17 | 0.52 | ns | VUS | Heterozygous germline. MSI-H endometrial tumor with loss of MSH2 and MSH6 by IHC. Two other clearly pathogenic loss-of-function *MSH2* mutations identified in the tumor make this less likely to be pathogenic | Y |
| Q374R | 0.20 | 0.42 | ns | B | Heterozygous germline. MSI-H endometrial tumor with loss of MLH1-MSH2, MSH6, and PMS2 by IHC. The tumor had 2 other clearly pathogenic somatic mutations in *MSH2*. | Y |
| V655I | 0.21 | 0.38 | ns | VUS | Heterozygous germline. Colorectal cancer under age 30. No family history of cancer. | N |
| W764R | 0.27 | 0.68 | ns | VUS | Heterozygous germline. MSI-H colon tumor with loss of MSH2 and MSH6 by IHC. Evidence of LOH for *MSH2* variant in tumor. | N |
| L599S | 0.33 | 0.39 | ns | VUS | Heterozygous germline. Seen in patient with breast cancer and family history of breast and colorectal cancer. | Y |
| V78I | 0.36 | 0.70 | ns | VUS | Heterozygous germline. MSS ovarian tumor with other pathogenic variants with no LOH for *MSH2* variant in tumor. | Y |
| I770V | 0.42 | 0.52 | ns | VUS | Heterozygous germline. Colorectal cancer diagnosed under age 30. Seen with heterozygous germline VUS in *APC* gene. | - |
| G761R | 0.47 | 0.38 | * | VUS | Suspected germline variant in prostate tumor. IHC for MSH2, MSH6 equivocal. MSI equivocal. Apparent *MSH2* LOH in tumor. | - |
| H785R | 0.75 | 0.51 | ns | VUS | Heterozygous germline. Colorectal cancer diagnosed over age 70. | - |
| P349A | 0.84 | 0.30 | *** | VUS | Homozygous germline. MSS colorectal cancer. The tumor also had *POLE* mutation and ultramutator phenotype. | - |
| Q681E | 1.06 | 0.48 | ** | VUS | Heterozygous germline. MSI-H colorectal cancer diagnosed over age 50. Tumor with loss of MLH1 and PMS2 explained by double somatic *MLH1* mutation in tumor. | N |
| A609P | 1.36 | 0.73 | ** | VUS | Heterozygous germline. MSI-H colon cancer with loss of MSH2 and MSH6 by IHC. Under age 50 at diagnosis. Tumor had one somatic pathogenic *MSH2* mutation seen at heterozygous frequency in the tumor. | Y |
| P696S | 1.71 | 0.98 | * | VUS | Heterozygous germline. Personal history of pheochromocytoma, family history of renal and brain cancer. | N |
| S676L | 2.06 | 0.70 | ** | VUS | Heterozygous germline. MSI-H colon cancer diagnosed under age 50. Tumor had loss of MSH6 by IHC. Seen with *MSH2* p.G827R somatic mutation listed below. | Y |

Table 3 (Continued)
| Human Genotype | Score<sup>a</sup> | 95% CI<sup>b</sup> | Sig<sup>c</sup> | ClinVar<sup>d</sup> | Clinical Information<sup>e</sup> | CST<sup>f</sup> |
| --- | --- | --- | --- | --- | --- | --- |
| C693R | 2.32 | 0.80 | **** | LP | Somatic mutation in tumor. MSI-H colon tumor with loss of MSH2 by IHC. This was seen with another heterozygous pathogenic mutation in *MSH2* (1760-1G>A). | Y |
| G692V | 2.42 | 0.58 | **** | LP | Somatic mutation in tumor. MSI-H neuroendocrine tumor with loss of MSH2 and MSH6 by IHC. Seen in a tumor with a germline likely pathogenic variant in *MSH2* (p.L30R) that had loss of heterozygosity. | - |
| G827R | 2.58 | 0.51 | **** | VUS | Somatic mutation in tumor. MSI-H colon cancer diagnosed under age 50. Tumor had loss of MSH6 by IHC. Seen with the germline variant p.S676L listed above. | Y |
| G692W | 2.66 | 0.61 | **** | VUS | Heterozygous germline. MSI-H endometrial tumor with loss of MSH2 and MSH6 by IHC. Cancer diagnosed over age 50. Variant reported by another laboratory to segregate with HNPCC in one family. A variant at the same position (p.G692R, NM_000179.2:c.2074 G to C) is classified as likely pathogenic (class 4) by the InSiGHT consortium. | Y |
| G827E | 2.80 | 0.63 | **** | VUS | Heterozygous germline. Pancreatic cancer diagnosed over age 80. Loss of *MSH2* and *MSH6* by IHC. | Y |
| N671D | 3.11 | 0.53 | **** | VUS | Heterozygous germline. Personal history of colon polyps and family history of colon, uterine, and other cancers. | Y |
| L310R | 3.25 | 0.74 | **** | P | Heterozygous germline. Seen in a family with multiple MSI-H colon cancers that had loss of *MSH2*. Co-segregation likelihood ratio for pathogenicity in the family was 44:1. See (Tsai et al., 2019) for complete pedigree information. | Y |

a. For stated genotype, variant mutation rate (mutations per generation) was compared to the WT mutation rate. The score represents Log<sub>2</sub>(median fold change).
b. 95% confidence interval on Log<sub>2</sub>(fold change).
c. Significance is calculated by a Wilcoxon Rank-Sum test with the Benjamini-Hochberg correction for multiple hypothesis testing. \* < 0.05 \*\* < 0.01 \*\*\* < 0.001 \*\*\*\* < 0.0001
d. Initial clinical classification as stated in ClinVar.
e. Clinical information collected from UW Laboratory Medicine. MSI-H = Microsatellite instability- high IHC = Immunohistochemistry
f. CST = Consistent, tumor or clinical data is consistent with functional score. Y = Yes, N = No, - = undetermined

Personal and family history of colorectal or endometrial cancer provide weak evidence of pathogenicity while personal and family history lacking HNPCC associated cancers provide weak evidence against pathogenicity (Li et al., 2020).Tumor characteristics of microsatellite instability (MSI-H) and loss of Msh2 and Msh6 on immunohistochemistry staining provide moderate evidence supporting pathogenicity (Li et al., 2020; Thompson et al., 2013). Presence of alternative pathogenic variants in *MSH2* or other genes that explain these tumor or other tumor characteristics provides evidence against pathogenicity, while a second somatic pathogenic variant at heterozygous frequency or loss of heterozygosity in tumor provides moderate and strong evidence supporting pathogenicity, respectively(Shirts et al., 2018). Formal strategies for combining each of these types of data with functional data are outside the scope of this effort. Rather we provide relevant clinical summaries with an overall assessment of whether the clinical information is consistent or inconsistent with functional scores. Of the 25 variants identified 16 (64%) had clinical data that was consistent with functional data, 4 (16%) had clinical data that was inconsistent with functional data, and 5 (20%) had clinical data that was equivocal or fell in the indeterminate functional score range. Some discordance is expected given that functional data is only one component of the ACMG guidelines for clinical variant interpretation; this discordance is consistent with results of past studies seeking to use other clinical criteria to classify variants(Li et al., 2020; Shirts et al., 2018; Thompson et al., 2013).

## Discussion

We developed a new method for high-throughput mutation rate assessment that combines a mid-20^th^ century method to determine mutation rate with 21^st^ century next generation sequencing. This allows for the pooling and multiplexing of mutation rate assessment which has not been accomplished before. We were able to determine the mutation rate of over 2,000 lineages representing ~200 variants of *MSH2*, a critical component of mismatch repair. Though we included a high frequency of WT sequences, our analysis indicates many of these could be substituted with additional VUS to increase throughput at minimal cost to accuracy. The assay is limited by the canavanine resistant subpopulation within a 200ml chemostat, which is dependent on the mutation rates of the lineages in a pool. One could increase the number of variants to be assayed in a single experiment by increasing the volume of the chemostat, though the logistics of expanding the volume beyond the 2L size of available commercial fermenters may be difficult. Another possible modification would be to utilize alternative marker loci that generate selectable mutations at higher rates than the wt *CAN1* sequence. In addition, the inclusion of barcoded null mutants may provide an additional control to better normalize the results to established mutation rates.

In this work, we estimated the pathogenicity of 157 variants of uncertain or conflicting significance derived from clinical testing. These results provide a key piece of information to testing labs seeking to assign pathogenicity to variants for which little other evidence is available. We have found that our results largely match clinical data obtained from tumors(Shirts et al., 2018). This data in combination with a recent deep mutation scan on Msh2 in a human cell line(Jia et al., 2020) will allow for accurate reclassification of uncertain and conflicting variants in this gene. This will provide physician’s better guidance on who to screen for HNPCC and how often.

This assay works with any protein which affects mutation rate, and so therefore can assess mutation rate variation in other proteins in the mismatch repair pathway, as well as in other DNA repair pathways. Assaying variants in the sequence context of the native human cDNA could be possible for genes that complement the orthologous yeast gene knockout(Kachroo et al., 2015; Vogelsang et al., 2009); however our initial attempts to recapitulate the complementation of *mlh1* and *pms1* with human *MLH1* and *PMS2*, other critical MMR genes associated with HNPCC, were unsuccessful. This, however, does not mean that assaying mutation rate of human alleles of DNA repair enzymes is not possible, and in fact would be a very interesting line of study.

The current set of alleles could also be tested in different genetic backgrounds, such as in a *san1Δ* background(Arlow et al., 2013) which could give additional information on the stability of Msh2 variants and could determine the mechanism behind an increased mutation rate for some variants. It could also be done in the background containing deletions or change of function mutations in other proteins in the MMR complex. Both variant library and background are completely mutable in this system.

Our method should be widely applicable and can be used to answer many other questions associated with mutation rate outside of clinical variant interpretation. Accurate, multiplexed measurement of mutation rate variation could be used to screen polymerases for increased or decreased fidelity, to screen the yeast deletion collection for knock-outs that increase mutation rate, or to uncover differences in mutation rate among natural variants of yeast. In conclusion, this method is broadly applicable to many different problems in which mutation rate is a factor and can be used to estimate the pathogenicity of clinically relevant DNA repair enzymes.

## Materials and Methods

### Growth in the chemostat

For individually inoculated chemostats, 1ml of overnight culture was inoculated into 230-245mls of glucose-limited media (Calcium Chloride 4g/L, Sodium Chloride 4g/L, Magnesium Sulfate Heptahydrate 20g/L, Potassium Phosphate Monohydrate 40g/L, Ammonium Sulfate 200g/L, Glucose 32g/L). Pools were thawed from the freezer and inoculated straight into the chemostat. Cells were allowed two days to grow to saturation and the pumps were turned on to a rate of 40mls/ hour or ~ 5 replacement volumes per day. Chemostats were allowed to reach steady state as determined by optical density and later confirmed by stabilization of CFU counts. Samples were then collected starting at 15 generations. Sampling the non-selective population involved spinning down ~ 2 × 10^8^ cells, as well as plating ~ 200 cells onto SC -histidine for accurate counts. For each chemostat, to determine the number of canavanine resistant mutants, sufficient culture to reach an estimated countable number of colonies was plated onto SC -arginine -serine -histidine + 60mg/L canavanine to select for loss-of-function mutations in *CAN1*. For pools, ~6 x 10^8^ cells were plated, in addition to those used for counts, onto 15cm SC + canavanine plates and then allowed to grow at 30C for 3 days at which point they were scraped for downstream analysis.

### Generation of Sequencing Libraries

For both the non-selective and mutant population, cells were vortexed vigorously with acid washed beads in resuspension buffer for 3 minutes, and then put through the Mini-prep Wizard kit. They were then concentrated using the PCR clean up Wizard kit.

For unbarcoded pools, *MSH2* was amplified from the plasmid vector using 15 rounds of PCR to prevent over-amplification. Then Nextera sequencing libraries were generated using the Nextera-XT kit. The average library size was 500bp, and sequenced using a Nextera 500 with paired end 150bp reads at a depth of ~30,000 reads over the length of *MSH2*. The run was conducted according to manufacturer specifications.

For barcoded pools: PCR amplification of the amplicon containing the barcode was done using 15-22 cycles of PCR using custom Nextera Primers, listed in Supplementary table 3. Sufficient amplification was determined by qPCR. The amplicon was purified using duel sided Sepharose bead cleanup to isolate the 250bp amplicon. Samples were then pooled at equal molar ratios and run on the NextSeq 550 using paired end reads of both 75 or 150 cycles using custom read and index primers (Supplementary table 3) at a read depth of ~1.7 million reads for the mutant population and 0.5 million reads for the non-selective population. Number of reads roughly corresponded with the number of colonies collected for the mutant pool, and 100X coverage of the known number of barcodes for the non-selective population.

### Data Analysis of Direct Sequencing of Unbarcoded Plasmid Pools

Reads were de-multiplexed using bcl2, allowing for no mismatches in the index read. Reads were then processed first by TrimGalore (FelixKrueger, 2019) to remove adaptors, then reads were collapsed using PEAR(Zhang et al., 2014) then aligned to the yeast *MSH2* sequence using BowTie2(Langmead and Salzberg, 2012), a SAM file was generated using Samtools(Li et al., 2009), and then the make up at each base pair was generated using Pysamstats (Miles, 2019). Data were manipulated in Excel, and then data points were graphed in R using ggplot(Wickham, 2009).

### Data Analysis Pipeline for Barcoded Pools

Reads were shortened to the barcode length using a custom python script, fed into PEAR to combine forward and reverse reads, then fed into Enrich(Fowler et al., 2011) to count barcodes. These counts were fed into a custom R script (supp file #1) which manipulated data and plotted using ggplot2.

### Competition of *msh2Δ* and WT

GFP was introduced to the *msh2Δ* and *his3Δ* strain by mating. Competitions were set up by individually inoculating 20ml glucose limited chemostats with 1ml of saturated culture of each stated strain. The strains were allowed to grow up for 2 days before the pumps were turned on. After reaching steady state after ~10 generations, cultures were mixed half and half, and GFP percentage was monitored twice a day via flow cytometry. Fitness effects were calculated by taking the slope of the ln of the GFP tagged to non-GFP tagged strains over time.

### Making of unbarcoded pools

DNA was extracted from *E. coli* strains sent from Alison Gammie and used to transform YMD4328: FY4 *msh2Δ* and *his3Δ* to ~20x coverage.

### Generating barcoded variants

Variants in human *MSH2* found in ClinVar were mapped to the yeast *MSH2* sequence. Variants containing a homolog within the yeast allele were then ordered as gene products from Twist Biosciences. The gene products were ligated into the pRS413 vector containing the yeast *MSH2* promoter and terminator and used to transform DH5α cells at 30X coverage. DNA was extracted using the Mira-Prep protocol (Pronobis et al., 2016) and then digested with SacI to linearize. A barcode along with randomized sequence were inserted into the linearized vector using Gibson assembly, and then used to transform DH5α cells. Transformants were collected such that there was 5X barcode coverage for each allele. DNA was extracted again using the Mira-prep protocol, and digested with SacI to linearize any unbarcoded alleles and transformed once more to take advantage of *E. coli’s* inability to be transformed by linear DNA that does not have homology overhangs. Colony PCR was done and 0% of clones contained no barcode and ~6% contained 2-3 barcodes. DNA was once again extracted with the Mira-Prep protocol, and then used to transform YMD4328: FY4 *msh2Δ* and *his3Δ flo1Δ* FY4. The *flo1Δ* is present to reduce the prevalence of flocs(Hope et al., 2017). Transformants were collected such that there was 20X coverage of each barcode. These were then pooled for future experiments.

### Pac-Bio Analysis

Plasmid fragments containing the barcode and variant were isolated from *E. coli* using the Wizard mini-prep kit, amplified using PCR with Kapa-HiFi, and cleaned up by digesting with DpnI and purifying with Ampure beads. Fragments were prepared for PacBio sequencing using the SMRTbell™ Template Prep Kit 1.0 (Pacific Biosciences) and sent to University of Washington PacBio Sequencing Services for sequencing and Sequel II circular consensus sequence (CCS) analysis(Wenger et al., 2019). BAM files of CCS reads were aligned to the plasmid reference using BWA/0.7.13 mem(Li, 2013). Reads that were aligned to the reference sequence were piped to a new BAM file with Samtools/1.9(Li et al., 2009) and analyzed with cigar strings to validate alignments. Barcodes were then extracted and two barcode-variant maps were generated. One file contains all the barcode-variant reads, and the other contains the highest quality read for each unique barcode. Errors found in these files were corrected using a multiple sequence alignment (MUSCLE 3.8.31)(Edgar, 2004) of reads sharing the same barcode. Final reads were derived from consensus sequences from these alignments. Ambiguous sequences were fixed by aligning sequences to the highest quality reads using the Needleman-Wunsch algorithm (EMBOSS 6.4.0)(Rice et al., 2000).

### Clinical Comparisons

Clinical comparisons were made using retrospective data gathered from clinical laboratory databases for testing performed as part of standard clinical care between 2014 and 2019. This retrospective analysis was done under University of Washington IRB 00007284.

### Data Availability

All sequencing data available at https://www.ncbi.nlm.nih.gov/bioproject/PRJNA662579. Strains and plasmids are available upon request. File S1 contains figures and tables referenced in this text, File S2 contains the custom scripts used in this work, File S3 contains strains and plasmids used in this text.

## Acknowledgements

We would like to thank Alison Gammie and Mark Rose for the strains used in this work. Josh Cuperus gave guidance on computational tools used in this project which we are grateful for. We appreciate Jolie Carlisle’s help with plasmid preps and initial attempts to develop this assay in an alternative system. We would like to thank Bryce Taylor, Pengyao Jiang, Jeet Patel, Meghan Garrett, Amanda Riley, Kurt Berckmueller, and Patrick Nugent for helpful edits on this manuscript. Our sincere gratitude to Jessica Hartmann for design help on figures. Research reported in this publication was supported by the National Cancer Institute of the National Institutes of Health (NIH) under award T32CA080416 (AWM), by the National Human Genome Research Institute of the NIH under award T32 HG00035 (ARO, AWM), and by the National Institute of General Medical Sciences of the NIH under award P41 GM103533 (MJD). MJD acknowledges prior support as a Rita Allen Foundation Scholar and as a Senior Fellow in the Genetic Networks program at the Canadian Institute for Advanced Research. This material is based in part upon work supported by the NSF under Cooperative Agreement No. DBI-0939454. The research of MJD was supported in part by a Faculty Scholar grant from the Howard Hughes Medical Institute.

